# TGF-β inhibitor SB431542 suppresses coronavirus replication through multistep inhibition

**DOI:** 10.1101/2025.03.20.644376

**Authors:** Assim Verma, Garvit Kumar, Nitin Khandelwal, Benjamin E. Mayer, Jitender Rathee, Yogesh Chander, Alka Nokhwal, Shweta Dhanda, Ram Kumar, Himanshu Kamboj, Riyesh Thachamvally, Shalini Sharma, Naveen Kumar

## Abstract

The COVID-19 pandemic highlighted the critical need for broad-spectrum antivirals with high resistance barriers. Here, we demonstrate that SB431542, a selective TGF-β receptor I (ALK5) inhibitor, exhibits potent antiviral activity against SARS-CoV-2 through unprecedented multitargeted mechanisms. Through comprehensive *in vitro*, and *in silico* analyses, we identified that SB431542 directly binds to SARS-CoV-2 ORF3a and disrupt its canonical function in inhibiting autophagosome-lysosome fusion. This interaction restored lysosomal acidification and normalized perinuclear LAMP-1 localization, significantly impairing virion assembly as evidenced by disrupted nucleocapsid-RNA association and reduced intracellular viral titers. Additionally, SB431542 downregulated the CLEAR network genes responsible for lysosomal biogenesis, further restricting viral egress pathways. Our temporal analyses revealed that at later infection stages (36-48 hpi), SARS-CoV-2 exploits TGF-β-induced lysosomal membrane permeabilization (LMP) and apoptosis for viral release—processes effectively inhibited by SB431542 through suppression of GADD45b and BAX expression. These multiple mechanisms resulted in an exceptional EC_50_ of 515 nM against SARS-CoV-2. *In vivo* efficacy was demonstrated in embryonated chicken eggs, where SB431542 conferred dose-dependent protection against lethal infectious bronchitis virus (IBV) challenge, with a favourable therapeutic index of 34.54. Remarkably, sequential passaging of SARS-CoV-2 for 50 generations under SB431542 selection pressure failed to generate resistant variants, contrasting sharply with the rapid resistance emergence typical of direct-acting antivirals. These findings establish SB431542 as a promising broad-spectrum coronavirus inhibitor with a unique triple-mechanism approach that simultaneously targets viral entry via TGF-β/Smad modulation, disrupts ORF3a-mediated lysosomal dysfunction affecting assembly, and attenuates TGF-β-induced apoptosis during late-stage infection— collectively imposing multiple selective constraints that impede escape mutation development.

**Importance:** The COVID-19 pandemic highlighted the urgent need for antiviral drugs with high barriers to resistance. This study reveals that SB431542, a drug previously developed to inhibit TGF-β signaling, exhibits remarkable effectiveness against SARS-CoV-2 through an unprecedented triple-mechanism approach. Unlike conventional antivirals that target a single viral component, SB431542 simultaneously disrupts viral entry, assembly, and release by binding to the viral ORF3a protein and modulating host cellular processes. Most importantly, SARS-CoV-2 failed to develop resistance against SB431542 even after 50 generations of exposure—a significant advantage over current therapeutics that quickly lose effectiveness due to viral mutations. Our findings also uncover that coronaviruses exploit both lysosomal dysfunction and programmed cell death to spread efficiently, providing new targets for therapeutic intervention. This research establishes SB431542 as a promising broad-spectrum coronavirus inhibitor and demonstrates the value of targeting host-virus interactions to overcome antiviral resistance.

## Introduction

The coronavirus disease 2019 (COVID-19) pandemic, caused by severe acute respiratory syndrome coronavirus 2 (SARS-CoV-2), precipitated an unprecedented global health crisis, necessitating the development of antiviral therapeutics with high barriers to resistance. SARS-CoV-2, infection cascade begins when the spike (S) glycoprotein engages the angiotensin-converting enzyme 2 (ACE2) receptor, followed by proteolytic priming via host proteases including TMPRSS2 and cathepsins, facilitating membrane fusion and viral genome release [1]. Unlike many enveloped viruses that utilize the conventional secretory pathway, SARS-CoV-2 distinctively exploits lysosomal exocytosis for virion release—a mechanism differentiating it from other coronaviruses [2, 3].

The viral accessory protein ORF3a serves as a master regulator of this lysosomal subversion process [2]. ORF3a inhibits autophagosome-lysosome fusion while simultaneously promoting lysosomal deacidification and exocytosis, thereby creating an environment conducive to virion preservation and efficient egress [2, 3].

This strategic manipulation of cellular degradation pathways not only facilitates viral dissemination but also compromises innate immune responses and cellular homeostasis [4, 5]. The centrality of lysosomal dysregulation in SARS-CoV-2 pathogenesis suggests that targeting these mechanisms could yield promising therapeutic outcomes [6, 7].

In addition to lysosomal dysregulation, SARS-CoV-2 induces profound alterations in host signaling networks, particularly the transforming growth factor-beta (TGF-β) pathway—a pleiotropic cascade that regulates autophagy, apoptosis, and inflammatory responses [8–13]. Hyperactivation of TGF-β/Smad signaling during SARS-CoV-2 infection enhances furin-mediated spike protein processing and cell-cell fusion events critical for viral propagation [8, 14]. Additionally, dysregulated TGF-β signaling contributes to the cytokine storms and pulmonary fibrosis characteristic of severe COVID-19 [15–17]. While elevated apoptotic indices have been documented in lung specimens from fatal COVID-19 cases, the mechanistic relationship between TGF-β-induced apoptosis and SARS-CoV-2 replication dynamics remains poorly studied [16, 18–20].

SB431542 is a potent and selective inhibitor of activin receptor-like kinase 5 (ALK5), a key mediator of TGF-β signalling [21]. In this study, we demonstrate that SB431542 exhibits antiviral potential against multiple coronaviruses. By using a combination of *in vitro*, *in silico*, and *in vivo* models, our findings reveal that beyond its canonical inhibition of TGF-β/Smad signaling, SB431542 exhibits previously unrecognized activity against SARS-CoV-2 through direct interaction with the viral ORF3a protein, thereby restoring lysosomal function.

Additionally, we uncovered a temporal dimension to SARS-CoV-2 egress mechanisms, wherein besides lysosomal exocytosis, the virus exploits apoptotic pathways during late-stage infection—a process effectively disrupted by SB431542. These mechanisms significantly reduce SARS-CoV-2-mediated cytopathic effects (CPE) in Vero cells while preventing IBV-induced stunted growth in embryonated chicken eggs.

A perennial challenge in antiviral therapeutics is the emergence of drug-resistant escape mutants under prolonged selection pressure [22, 23]. Resistance mutations often compromise drug efficacy and limit long-term therapeutic utility. Besides antiviral mechanistic insights, we also addressed this issue by subjecting SARS-CoV-2 to 50 serial passages under SB431542 pressure. Notably, no resistant mutants were observed, underscoring SB431542’s high genetic barrier to resistance.

Collectively, our findings highlight SB431542 as a potential therapeutic candidate with multiple antiviral mechanisms—targeting both TGF-β signalling and ORF3a-mediated lysosomal dysfunction—with a remarkably high genetic barrier to resistance.

## Results

### SB431542 suppresses SARS-CoV-2 entry and release from Vero cells during the early hours of infection

In a comprehensive screening of approximately 150 host kinase inhibitors, we identified the transforming growth factor-β (TGF-β) receptor I (ALK5) antagonist SB431542 as a potent inhibitor of SARS-CoV-2 replication. SB431542 exhibited dose-dependent antiviral efficacy in Vero cells at non-cytotoxic concentrations **(Fig. 1A)**, with an EC_50_ of 515 nM **(Fig. 1B)**. This potency substantially exceeds that of the FDA-approved nucleoside analog remdesivir (EC_50_: 1.65 μM) in comparable cellular systems [24], positioning SB431542 as a potent inhibitor against SARS-CoV-2.

**Figure 1.**
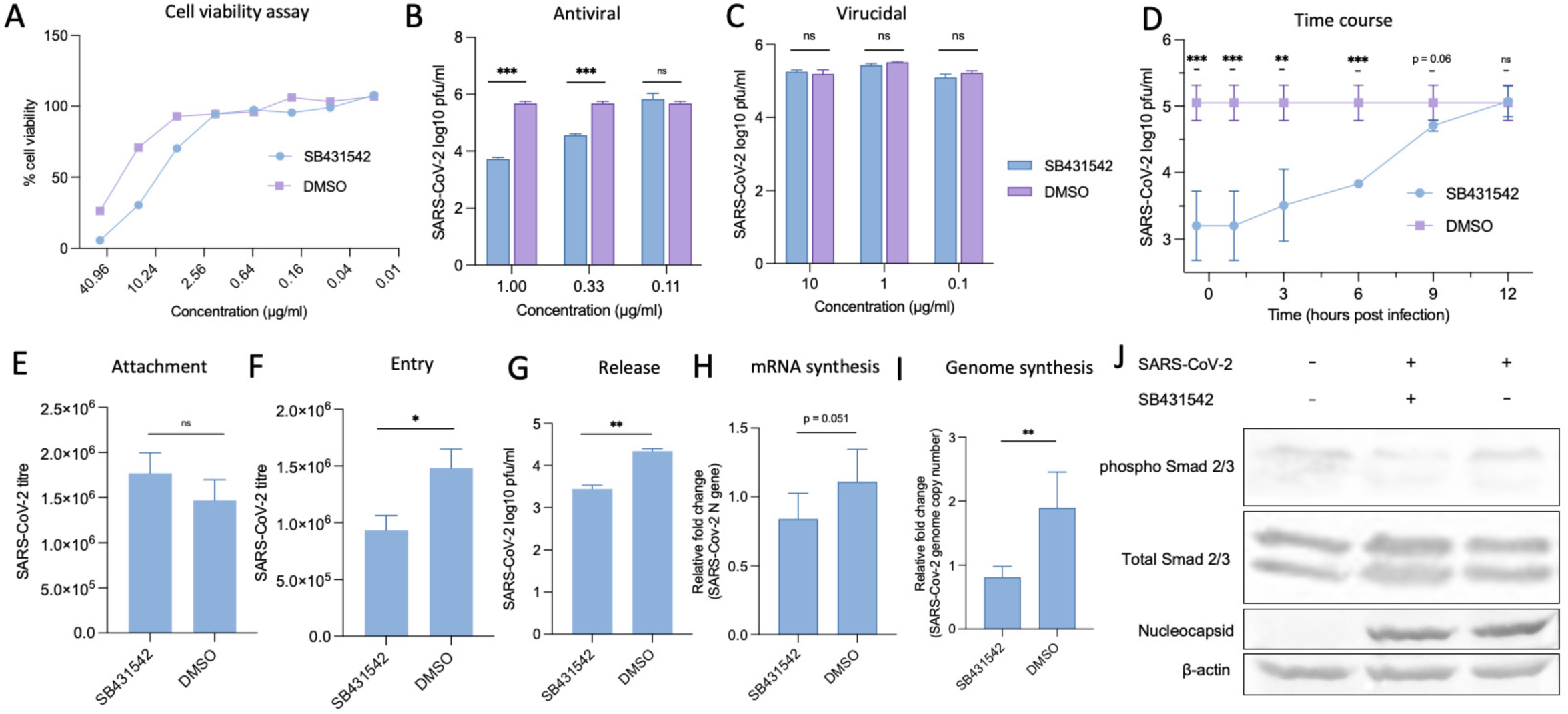
SB431542 treatment suppresses SARS-CoV-2 replication. **A. Cytotoxicity of SB431542 (MTT assay):** Vero cells were treated with the indicated concentrations of SB431542 or equivalent volumes of DMSO in triplicates for 96 hours. Cell viability was measured using the MTT assay, and the CC50 was calculated using the Reed-Muench method. **B. *In vitro* anti-SARS-CoV-2 efficacy:** Vero cells were infected with SARS-CoV-2 at an MOI of 0.1 and treated with indicated concentration of SB431542 or DMSO in triplicates. Infectious virus particles in the culture supernatants were quantified at 72 hpi by plaque. **C. Virucidal activity:** SARS-CoV-2 was incubated with the indicated concentrations of SB431542 or equivalent volumes of DMSO for 90 minutes at 37°C. The virus-inhibitor mixture was diluted (1:1000), and residual viral infectivity was determined by plaque assay **D. Time-of-addition assay:** Confluent monolayers of Vero cells were infected with SB431542 at an MOI of 5, washed with PBS, and treated with SB431542 or DMSO at indicated timepoints. Supernatants were collected at 12 hpi, and infectious virions were quantified by plaque assay. **E. Attachment assay:** Vero cells, in triplicates, were preincubated with SB431542 or DMSO for 1 hour before infection with SARS-CoV-2 at an MOI of 5 for 1 hour at 4°C to allow attachment. Cells were then washed five times with PBS, lysed by rapid freeze-thaw, and the virus released was quantified by plaque assay. **F. Entry assay:** Vero cells, in triplicates, were infected with SARS-CoV-2 at an MOI of 5 for 1 hour at 4°C in SB431542-free medium to allow attachment. After washing five times with ice-cold PBS, fresh medium containing SB431542 or DMSO was added, and cells were incubated at 37°C for 1 hour to permit viral entry. Cells were subsequently washed and incubated with SB431542-free DMEM. Virus particles released at 12 hpi were titrated by plaque assay. **G. Virus release assay:** Vero cells, in triplicates, were infected with SARS-CoV-2 at an MOI of 5 for 1 hour, washed five times with PBS, and supplemented with fresh DMEM. At 8 hpi, when virus budding presumably begins, cells were washed again and fresh medium containing SB431542 or DMSO was added. Virus particles released into the supernatant at 4 hours post-drug treatment were quantified by plaque assay. **H, I. Viral mRNA/genome synthesis:** Vero cells were infected with SARS-CoV-2 at an MOI of 5, washed with PBS, and treated with SB431542 or DMSO at 2 hpi. RNA and DNA were isolated from cells harvested at 6 hpi. For mRNA quantification, cDNA was synthesized from RNA using oligo(dT) primers. SARS-CoV-2 *N* gene expression was quantified by qRT-PCR, normalized to β-actin (housekeeping control), and analyzed using the ΔΔCt method. **g. Western blot analysis:** Vero cells were infected with SARS-CoV-2 and treated with SB431542 or vehicle control at 2 hpi. At 12 hpi, cell lysates were prepared in RIPA buffer and analyzed by Western blot using anti-SARS-CoV-2 Nucleocapsid (upper panel) or GAPDH antibody (lower panel). Relative band intensities were quantified using ImageJ software and normalized to GAPDH as a loading control (bottom). All values represent means ± SD from at least three independent experiments. Statistical significance was determined using Student’s t-test (ns = non-significant, *P < 0.05; **P < 0.01; ***P < 0.001).

To delineate whether SB431542 directly neutralizes extracellular virus, we conducted virucidal assays wherein SARS-CoV-2 was pre-incubated with SB431542 at concentrations up to 10-fold above the non-cytotoxic threshold. The absence of reduced infectivity **(Fig. 1C)** indicated that SB431542 does not compromise virion integrity directly but likely targets intracellular replication processes.

To further elucidate the specific stages of the SARS-CoV-2 life cycle affected by SB431542, we performed a time-of-addition assay by infecting Vero cells at an MOI of 5, and adding the inhibitor at various time points post-infection. Addition of SB431542 either 30 minutes pre-infection or within the first hour post-infection (hpi) conferred maximal inhibition, with a gradual attenuation of efficacy when added at later timepoints **(Fig. 1D)**. This temporal sensitivity pattern suggested interference with both early viral entry mechanisms and late-stage post-replicative processes.

To precisely elucidate the specific phases of the viral lifecycle targeted by SB431542, we performed stage-specific assays. Temperature-restricted (4°C) attachment experiments demonstrated that SB431542 does not significantly impair the initial virion-receptor engagement **(Fig. 1E)**. However, when pre-attached virions were permitted to internalize by temperature elevation to 37°C, SB431542 treatment significantly suppressed productive infection **(Fig. 1F)**, indicating disruption of post-attachment entry mechanisms. Additionally, when administered at 12 hpi following the removal of pre-released virions by extensive washing, SB431542 significantly reduced subsequent virion egress **(Fig. 1G)**, establishing its capacity to inhibit late-stage virion release.

Importantly, when SB431542 was applied during the intermediate phase (3hpi), neither intracellular viral RNA accumulation **(Fig. 1H, I)** nor viral protein synthesis **(Fig. 1J)** was appreciably affected, while phosphorylation of Smad2/3, the canonical mediator of TGF-β signalling was markedly attenuated [25, 26].

Taken together, these results indicate that SB431542 inhibits SARS-CoV-2 entry and release without affecting other steps such as RNA or protein synthesis. While its effect on viral entry has been previously reported [8], our findings highlight its additional role in disrupting egress mechanisms.

### SB431542 restore lysosomal function by subverting ORF3a

SARS-CoV-2 uniquely exploits lysosomal exocytosis for viral egress, a process orchestrated by its accessory protein ORF3a [2, 3]. Unlike its counterpart in SARS-CoV, the SARS-CoV-2 ORF3a protein inhibits autophagosome-lysosome fusion while simultaneously promoting lysosomal deacidification, thereby creating an environment conducive to virion preservation and efficient release [2, 27]. To investigate whether SB431542 targets this pathway, we examined autophagosome dynamics using GFP-LC3 as a marker for autophagosome accumulation [28].

Consistent with previous studies, overexpression of GFP-LC3 followed by SARS-CoV-2 infection led to cytoplasmic foci formation indicative of autophagosome accumulation [5, 29] **(Fig. 2A)**. However, treatment with SB431542 inhibited this aggregation and dispersed GFP-LC3 throughout the cytoplasm. This effect was further validated in HeLa cells overexpressing ORF3a and GFP-LC3 wherein, treatment with SB431542 significantly reduced LC3-GFP puncta formation **(Fig 2B).** These observations suggest that SB431542 interferes with ORF3a-mediated disruption of autophagosome-lysosome fusion. It is noteworthy that TGF-β1 signalling is well-established in autophagic induction [15, 30], the apparent autophagy-inducing effect of its putative inhibitor, SB431542, in the context of SARS-CoV-2 infection was intriguing.

**Figure 2.**
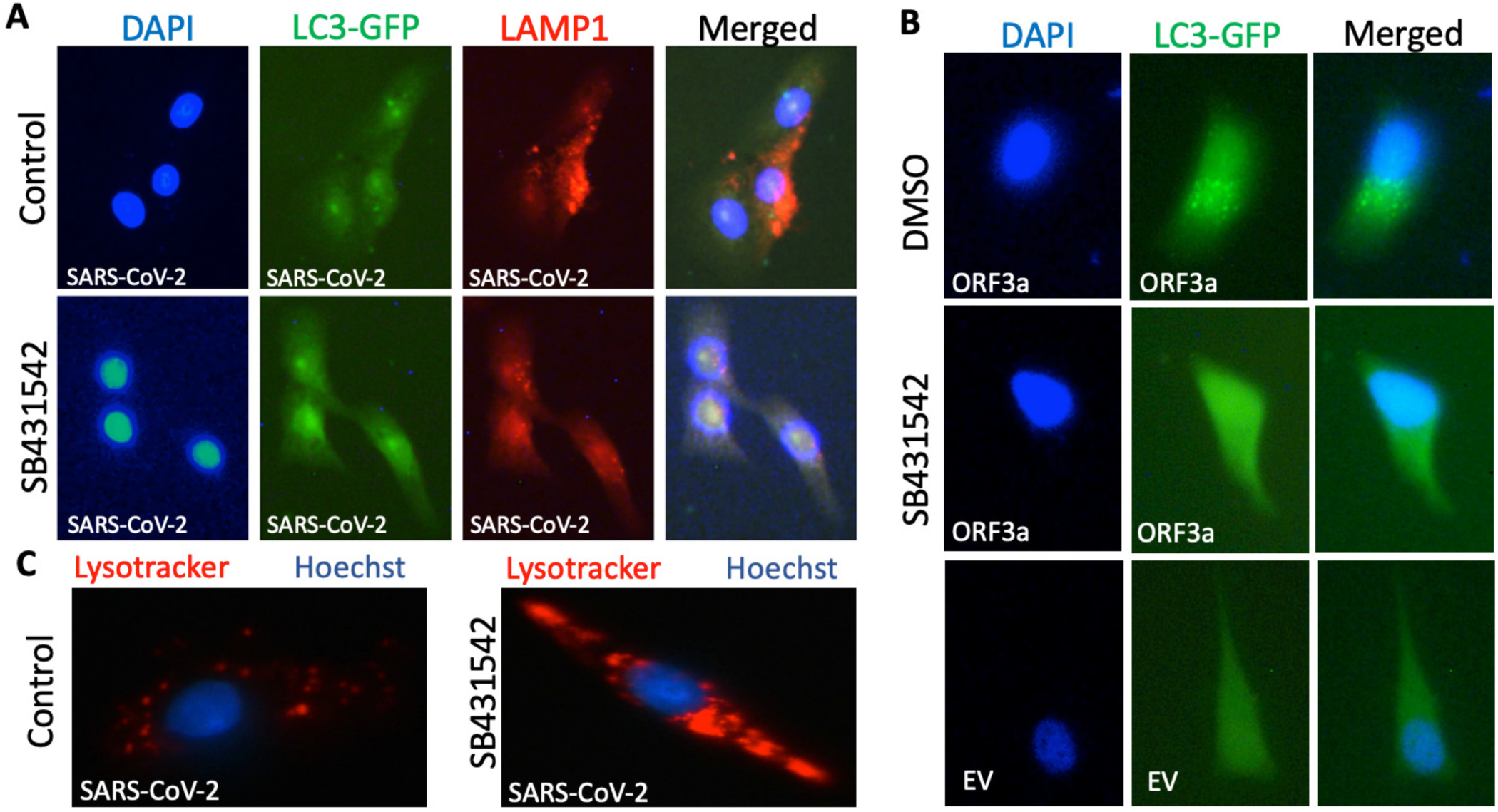
SB431542 disrupts ORF3a-mediated lysosomal dysregulation. **Immunofluorescence imaging of A. Vero cells transfected with LC3-GFP followed by SARS-CoV-2 infection:** Vero cells were transfected with LC3-GFP. After 48 hours SARS-CoV-2 was inoculated at an MOI of 5 for 1 hour, followed by washing with PBS and incubation for 12 hours with SB431542 (lower panel) or DMSO (upper panel). At 12 hpi, cells were fixed and permeabilized followed by probing with anti-LAMP1 antibody and DAPI . **B. HeLa cells transfected with LC3-GFP :** HeLa cells were transfected with LC3-GFP and SARS-CoV-2 ORF3a or empty vector for 48 hours and treated with SB431542 or vehicle control, followed by fixing and permeabilization. DAPI was used to stain nucleus. **C. Lysotracker assay:** Vero cells were infected with SARS-CoV-2 at 1 MOI for 1hour followed by treatment with DMSO (left panel) or SB431542 (right panel). At 24 hpi cells were stained with Lysotracker and Hoechst dyes and analysed with fluorescent microscope.

### SB431542 targets SARS-CoV-2 ORF3a at the saposin-a interaction interface

To elucidate the potential mechanism underlying this phenomenon, we conducted a literature survey and found that ORF3a is a potential target of macro-heterocyclic compounds such as chlorin [31]. This prompted us to perform in-depth computational analyses including protein-ligand structural modelling and molecular dynamics simulations.

Using the diffusion-based structure prediction tool Boltz [32], we generated protein-ligand complexes based on the resolved ORF3a dimer structure (PDB ID: 6XDC). The modelling revealed that SB431542 binds to a specific cavity within the ORF3a dimer **(Figure 3A)**. As a comparative control, we modelled the binding of bacteriochlorin, a structural analog of chlorin previously reported to interact with ORF3a **(Figure 3B)** [31]. Detailed examination via Ligplot analysis [33] of the protein-ligand interfaces demonstrated that both compounds established interactions with similar residues in the ORF3a dimer **(Fig. 3C, D)**.

**Figure 3.**
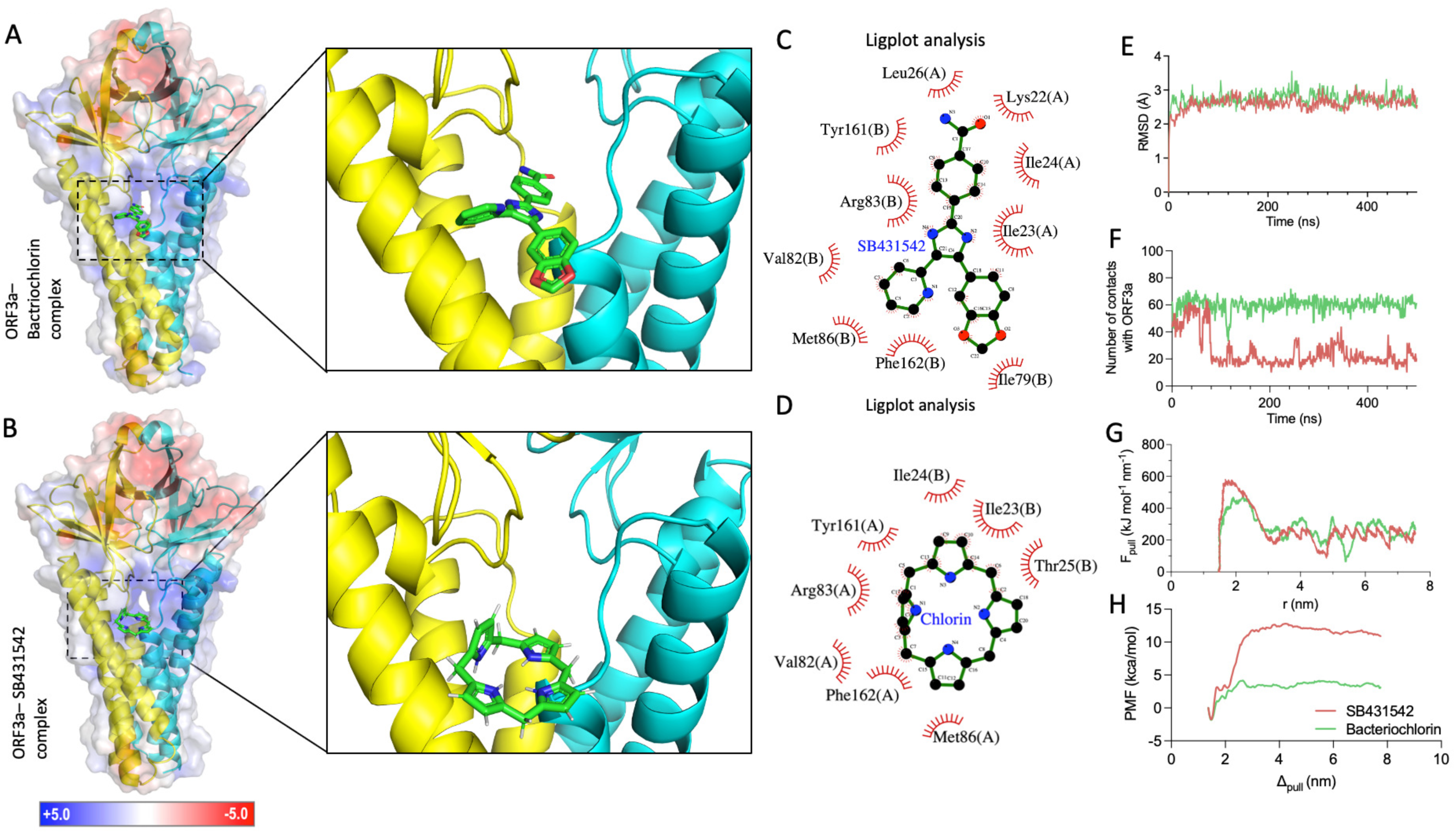
SB431542 binds to ORF3a via off-target effect. **A. ORF3a – SB431542 complex (Boltz prediction):** An illustration of the Boltz generated ORF3a – SB431542 structure is shown. The ORF3a chains are colored in yellow and cyan and color coded surface plot of the electrostatics is shown as well. The side shows a zoom into the SB431542 binding region. **B. ORF3a – Bacteriochlorin complex (Boltz prediction):** An illustration of the Boltz generated ORF3a – Bacteriochlorin structure is shown. The ORF3a chains are colored in yellow and cyan and color coded surface plot of the electrostatics is shown as well. The side shows a zoom into the Bacteriochlorin binding region. **C, D. ORF3a – ligand interactions:** Liglot analysis of SB431542 **(C)** and bacteriochlorin **(D)** bound to ORF3a. **E. RMSD of protein complexes:** The root mean square fluctuation of the ORF3a complexes from both performed 500ns MD simulations is plotted over the simulated time range. Though fluctuations are visible a general stabilization between 2Å and 3Å is visible. **F. Number of protein-ligand contacts over time:** Here the absolute number of contacts based on a 4Å cutoff between the ligand and ORF3a dimer is illustrated over simulated time. While the number of contacts oscillate at around 60 contacts for Bacteriocholorin a drop to around 20 contacts can is clearly visible at 100ns. **G. Non-equilibrium pulling force:** The pulling force observed in the non equilibrium pulling simulation is illustrated of simulation time for both ORF3a-SB431542 and ORFa-Bacteriochlorin complexes. A overall higher peak of to be overcome force can be observed for SB431542 (583 kJ/mo/nm) compared to Bacteriochlorin (470 kJ/mol/nm). **H**. **Potential of mean forces profile:** The two generated potential of mean forces profiles are illustrated over pulling distance for both investigated ligands. After a first decrease in PMF both curves continuously increase until reaching a plateau at around 4 nm. While the absolute PMF value shows a slight downward trend from 4 nm on the absolute difference towards the minimal PMF value is significantly higher, implying a stronger ΔG value.

Molecular dynamics simulations (500 ns) demonstrated remarkable stability of both protein-ligand complexes, as evidenced by minimal RMSD fluctuations throughout the simulation period **(Fig. 3E)**. Additional analyses including RMSF and radius of gyration (Rg) further confirmed the structural integrity of the complexes during simulation **(Supplementary Fig. 1A, B)**. Quantitative assessment of protein-ligand interactions revealed distinct binding characteristics. While bacteriochlorin maintained approximately 60 contacts with ORF3a throughout the simulation, SB431542 exhibited fewer contacts (approximately 20-30) during equilibrium phases **(Fig. 3F)**. Despite this difference in contact profiles, both compounds adopted similar binding modes within the ORF3a structure **(Supplementary Fig. 1C)**.

To understand the interaction dynamics and binding affinity we first performed MM-GBSA analysis [34, 35], which revealed slightly lower average interaction energy for ORF3a-SB431542 compared to ORF3a-bacteriochlorin **(Supplementary Figure 1D)**. This prompted us to perform non-equilibrium pulling steered molecular dynamics (SMD) which demonstrated that SB431542 required significantly greater force for extraction from its binding pocket compared to bacteriochlorin **(Fig. 3G)**, suggesting stronger binding interactions despite maintaining fewer contacts.

The more rigorous umbrella sampling approach further provided definitive evidence of the superior binding affinity of SB431542 [36]. Analysis of the potential of mean force (PMF) between protein and ligand, illustrated in **Figure 3H**, revealed substantially stronger energetic differences for SB431542 compared to bacteriochlorin. Calculating all possible differences and using the mean values **(Supplementary Figure 1E)**, we determined binding free energies of ΔG = −13.32 kcal/mol for SB431542 and ΔG = −5.28 kcal/mol for bacteriochlorin.

Intriguingly, structural alignment with the recently published ORF3a-Saposin-A complex (PDB ID: 8EQU) revealed that the SB431542 binding site directly overlaps with the protein-protein interaction interface between ORF3a and Saposin-A [37] **(Supplementary Fig. 1F)**. Higher magnification views of both the initial and final states of the MD simulation clearly demonstrated that SB431542 occupies the precise interface where critical interactions between ORF3a and Saposin-A occur **(Supplementary Fig. 1G, 1H)**. This structural insight provides a mechanistic explanation for the ability of SB431542 to inhibit lysosomal exocytosis pathways. Together, these computational findings corroborate our experimental observations that SB431542 disrupts ORF3a-mediated lysosomal dysfunction, thereby inhibiting SARS-CoV-2 egress.

### SB431542 treatment restores lysosomal pH

Besides LC3, we also examined lysosomal function by analyzing LAMP-1 localization, a key lysosomal membrane protein involved in lysosomal biogenesis, autophagy, and SARS-CoV-2 exocytosis [2, 38, 39]. In SARS-CoV-2-infected cells, LAMP-1 predominantly localized to peripheral regions indicative of active lysosomal exocytosis [2] **(Fig. 2A)**. In contrast, SB431542 treatment sequestered LAMP-1 within perinuclear regions. Additionally, SB431542-treated cells exhibited lower LAMP-1 expression compared to vehicle controls, suggesting inhibition of the canonical ORF3a function that typically increases LAMP-1 expression following SARS-CoV-2 infection [2, 27].

Given that, peripheral lysosomes are less acidic than juxtanuclear ones [40], we tracked the lysosomes using lysotracker dye which give proximation of both location and pH of lysosomes. As expected, SARS-CoV-2 infected cells displayed peripheral lysosome localization with diminished fluorescence intensity, indicative of deacidification and active exocytosis **(Fig. 2C).** Treatment with SB431542 restored fluorescence intensity and redistributed lysosomes to perinuclear regions, indicating restoration of lysosomal acidification. These evidences collectively suggest that SB431542 treatment restored lysosomal function by targeting ORF3a.

### SB431542 disrupts SARS-CoV-2 assembly

To investigate the functional consequences of SB431542-mediated restoration of lysosomal function on SARS-CoV-2 egress, we performed a several morphological and biochemical analyses. Bright-field microscopy revealed pronounced differences in cellular architecture between control and SB431542-treated infected cells at 12 hpi. Control-infected cells exhibited substantial lysosomal vacuolization **(Fig. 4A, upper panel)**, with particularly prominent vacuolar structures visible under phase contrast **(Fig. 4B, upper panel)**.

**Figure 4.**
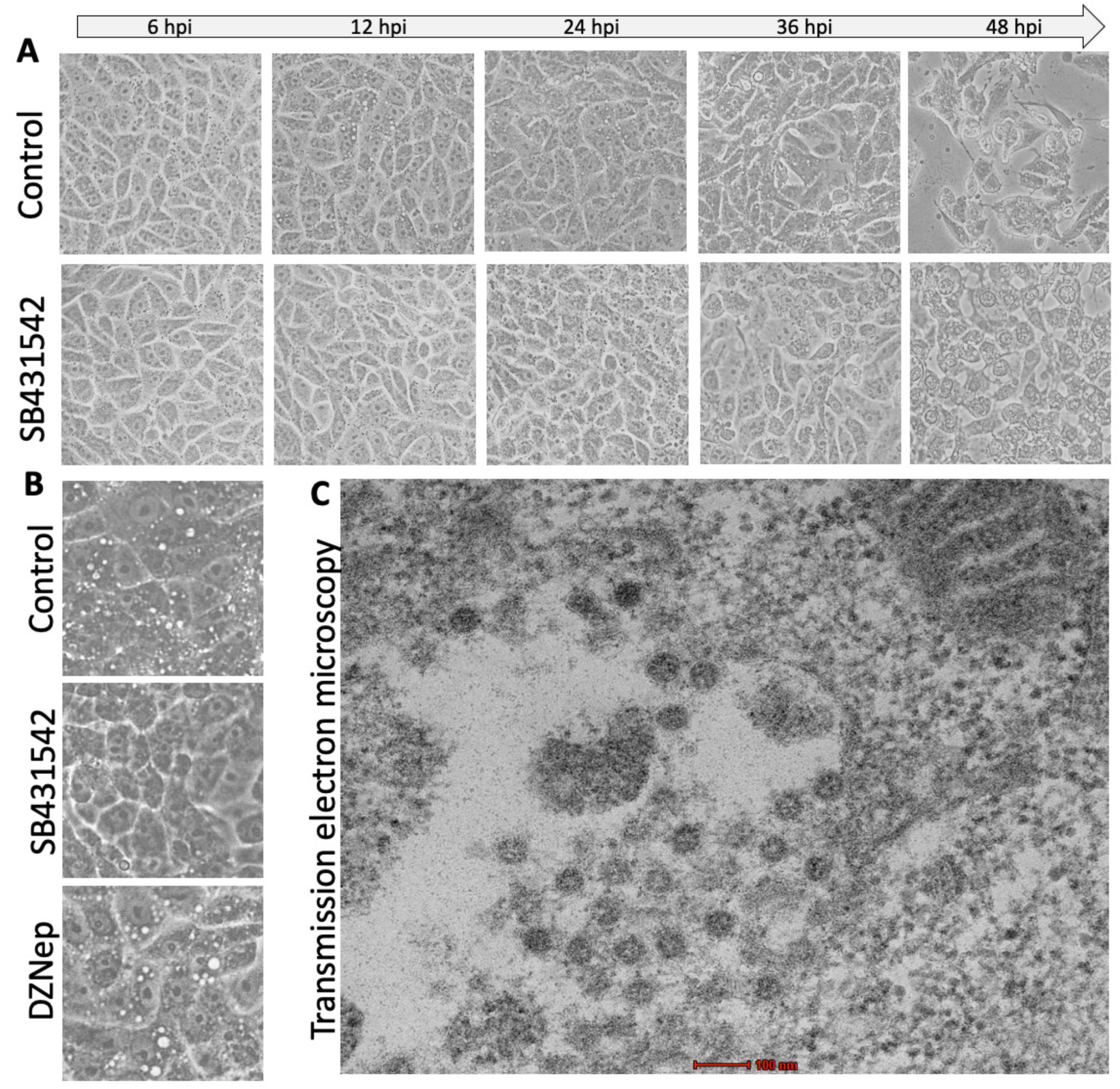
SB431542 disrupts SARS-CoV-2 mediated lysosomal vacuolization. **A. Bright field microscopy:** Vero cells were infected with SARS-CoV-2 at 5 MOI for 1 hour followed by treatment with DMSO (upper panel) or SB431542 (lower panel). Images were taken at indicated time-points. **B. Phase contrast microscopy:** Vero cells were infected with SARS-CoV-2 at 5 MOI for 1 hour followed by treatment with DMSO (upper panel), SB431542 (middle panel) or DZNep (lower panel). Images were taken at 12 hpi. **C. Transmission electron microscopy:** Vero cells were infected with SARS-CoV-2 at 5 MOI for 1hour. At 12 hpi cells were fixed using glutaraldehyde solution and subsequently blocks were prepared and thin sectioning was performed for TEM imaging.

SB431542 treatment remarkably suppressed this vacuolization phenotype **(Fig. 4A, lower panel; Fig. 4B, middle panel)**. We also examined the effect of DZNep, another SARS-CoV-2 inhibitor with a distinct mechanism of action [41], which failed to prevent vacuolization **(Fig. 4B, lower panel)**, indicating the specificity of the action of SB431542. Transmission electron microscopy (TEM) analysis further confirmed that these vacuoles contained assembled SARS-CoV-2 virions **(Fig. 4C)**, implicating lysosomal dysfunction and subsequent vacuolization in viral assembly and release [38].

To quantitatively assess the impact of SB431542 on virion assembly and egress, we conducted parallel analyses of intracellular and extracellular viral loads. Following infection at MOI 1, SB431542 treatment significantly reduced both intracellular virion abundance **(Fig. 5A)** and extracellular viral titers **(Fig. 5B)** at 24 hpi. This concurrent diminution of both intracellular and extracellular viral populations suggested that SB431542 disrupts virion biogenesis upstream of the egress process. Combined with data from **Fig. 2**, these results suggest that SB431542-mediated lysosomal acidification disrupts the production of infectious virions.

**Figure 5.**
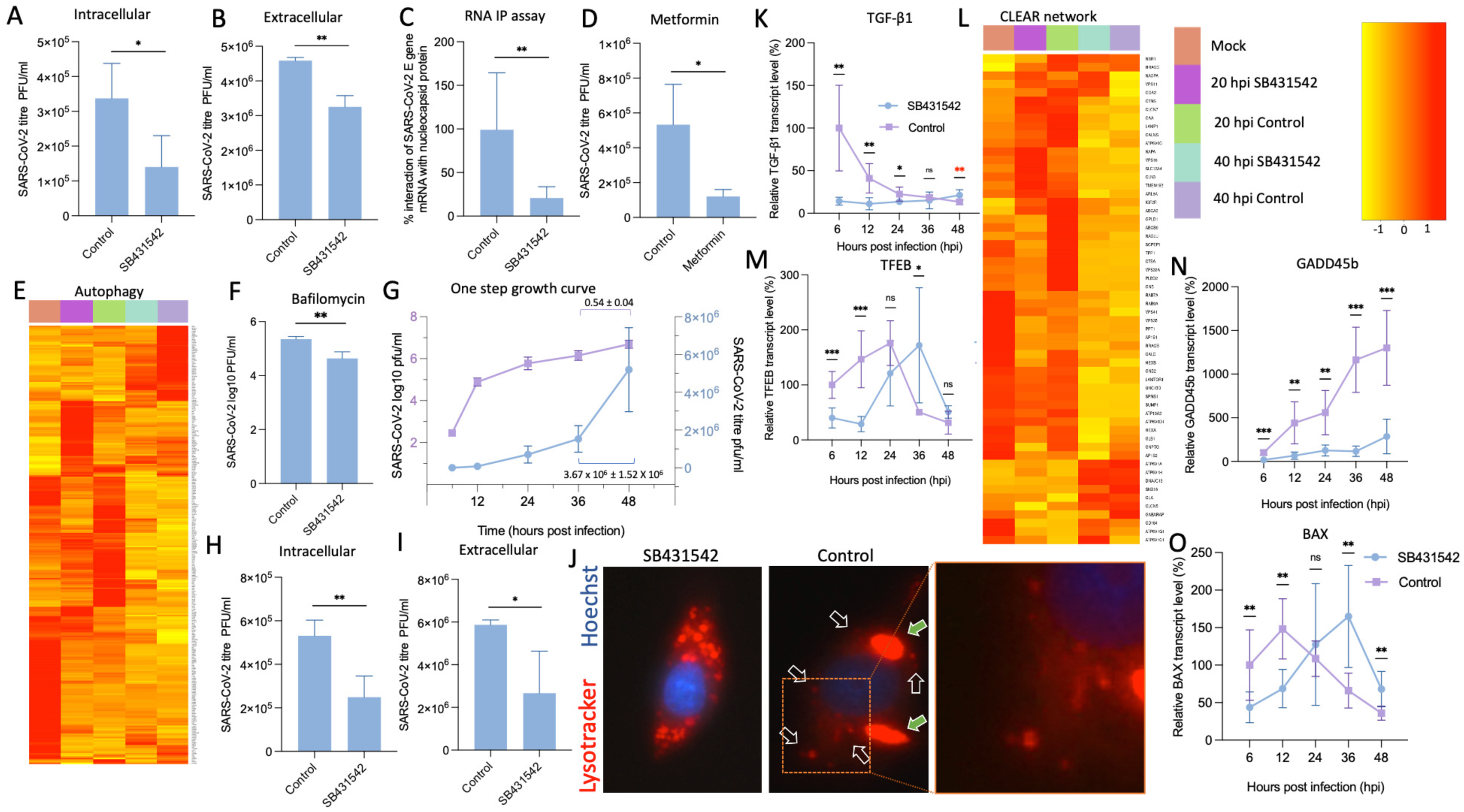
SB431542 blocks SARS-CoV-2 release. **A, B Quantification of intra- and extracellular virions at 24 hpi:** Vero cells were infected with SARS-CoV-2 at 1 MOI for 1 hour followed by treatment with DMSO or SB431542. At 24 hpi supernatants were harvested and cells were washed with PBS multiple times, followed by rapid freeze-thaw cycles to retrieve intracellular virions. Infectious virions from supernatant and cell lysates were quantified with plaque assay. **C. RNA-IP assay:** Vero cells were infected with SARS-CoV-2 at an MOI of 5 and treated with SB431542 or DMSO at 2 hpi. At 16 hpi, cell lysates were prepared for RNA-IP as described in Materials and Methods. Lysates were incubated with α-nucleocapsid (reactive antibody), α-ERK (non-reactive antibody), or IP buffer alone (beads control), followed by Protein A Sepharose slurry incubation. After washing and cross-link reversal, cDNA was from immunoprecipitated RNA and SARS-CoV-2 RNA (N gene) was quantified by qRT-PCR and normalized to input controls. **D. Metformin antiviral assay:** Vero cells were infected with SARS-CoV-2 at 0.1 MOI for 1 hour followed by treatment with metformin or vehicle control. At 24 hpi supernatants were harvested and virus were quantified using plaque assay. **E. RNAseq analysis of autophagy related genes:** Vero cells were infected with SARS-CoV-2 at 5 MOI for 1 hour followed by treatment with DMSO or SB431542. At 12 hpi, cells were scraped and total RNA was isolated followed by RNAseq analysis as mentioned in methods section. For analysis of autophagy related genes from samples, KEGG pathway filtering was utilized. **F Bafilomycin antiviral assay:** Vero cells were infected with SARS-CoV-2 at 0.1 MOI for 1 hour followed by treatment with bafilomycin or vehicle control. At 24 hpi supernatants were harvested and virus were quantified using plaque assay. **G. One-step growth curve analysis.** Vero cells were infected with SARS-CoV-2 at 1 MOI for 1 hour followed by washing with PBS and supplemented with fresh DMEM. Supernatants were harvested at indicated timepoints and virus were quantified using plaque assay. **H, I Quantification of intra- and extracellular virions at 48 hpi:** Vero cells were infected with SARS-CoV-2 at 1 MOI for 1 hour followed by treatment with DMSO or SB431542. At 48 hpi supernatants were harvested and cells were washed with PBS multiple times, followed by rapid freeze-thaw cycles to retrieve intracellular virions. Infectious virions from supernatant and cell lysates were quantified with plaque assay. **J. Lysotracker assay:** Vero cells were infected with SARS-CoV-2 at 1 MOI for 1hour followed by treatment with DMSO (left panel) or SB431542 (right panel). At 42 hpi cells were stained with Lysotracker and Hoechst dyes and analysed with fluorescent microscope. **K. Quantification of TGF-β1 gene:** Vero cells were infected with SARS-CoV-2 at 1 MOI for 1 hour followed by treatment with DMSO or SB431542. At indicated time-points, cells were scraped and RNA was isolated, followed by cDNA library preparation using oligo(dT). TGF-β1 genes were amplified from the harvested cells and normalized to β-actin gene (housekeeping control). Relative fold-change was calculated using the ΔΔCt method. **L. RNAseq analysis of CLEAR network related genes:** Vero cells were infected with SARS-CoV-2 at 5 MOI for 1 hour followed by treatment with DMSO or SB431542. At 12 hpi, cells were scraped and total RNA was isolated followed by RNAseq analysis as mentioned in methods section. For analysis of CLEAR network related genes from samples, KEGG pathway filtering was utilized**. M, N and O. Quantification of GADD45b, TFEB and BAX genes:** Vero cells were infected with SARS-CoV-2 at 1 MOI for 1 hour followed by treatment with DMSO or SB431542. At indicated time-points, cells were scraped and RNA was isolated, followed by cDNA library preparation using oligo(dT). GADD45b **(M),** TFEB **(N)** and **(O)** BAX genes were amplified from the harvested cells and normalized to β-actin gene (housekeeping control). Relative fold-change was calculated using the ΔΔCt method. Values represent means ± SD from at least three independent experiments. Statistical significance was determined using Student’s t-test (ns = non-significant, *P < 0.05; **P < 0.01; ***P < 0.001).

The nucleocapsid (N) protein plays a central role in coronavirus assembly by encapsidating viral genomic RNA [42, 43]. RNA immunoprecipitation assays using anti-N antibodies revealed that SB431542 treatment significantly abrogated the association between N protein and viral RNA **(Fig. 5C)**, indicating disruption of a critical initial step in virion assembly.

Given the established role of lysosomal acidification in regulating autophagic processes [44], we hypothesized that the effect of SB431542 on viral assembly might be mechanistically linked to the restoration of autophagic flux. To test this, we employed metformin, an over the counter available drug known for autophagy inducer [45] which significantly reduced SARS-CoV-2 titers by 72.5% with EC_50_ ∼35 µM at non-cytotoxic dose of 50 µM **(Fig. 5D)**, phenocopying the effect of SB431542. Transcriptomic analysis further revealed that SB431542 modulated the expression of autophagy-related genes at both early (20 hpi) and late (40 hpi) infection timepoints **(Fig. 5E).** Conversely, complete inhibition of autophagy using bafilomycin A1 also suppressed viral titers **(Fig. 5F)**, consistent with previous studies demonstrating that SARS-CoV-2 requires induction of early autophagy while inhibiting later stages of autophagosome-lysosome fusion [46]. Taken together, these findings demonstrate that SB431542 restores lysosomal acidification and autophagic flux by targeting ORF3a, thereby disrupting SARS-CoV-2 assembly.

### SB431542 suppresses SARS-CoV-2 infection through canonical inhibition of apoptosis

While SARS-CoV-2 has a replication cycle of approximately 8 h [41, 47, 48], significant cytopathic effects (CPE) were only observed at 42–48 hpi even at high MOI of 5 **(Fig. 4A)**. Prior to this time point, infected cell morphology remained comparable to uninfected controls, consistent with previous observations that viral egress does not compromise cell viability until later stages of infection [49]. This observation is particularly noteworthy as most previous studies investigating SARS-CoV-2 egress mechanisms were temporally restricted to early infection stages (16-24 hpi), during which lysosomal exocytosis serves as the primary release mechanism [2, 49, 50].

To characterize viral replication dynamics during this extended infection period, we performed one-step growth curve analysis up to the onset of observable CPE. Between 36 and 48 hpi, we documented a substantial 340% increase in viral titers, equivalent to a linear-scale increment of 3.67×10^6^ ± 1.52×10^6^ **(Fig. 5G)**. SB431542 treatment significantly suppressed both intracellular and extracellular viral titers even at 48 hpi **(Fig. 5H, 5I).**

Lysotracker staining at 42 hpi, a timepoint where CPE was starting to occur, revealed stress-induced lysosome membrane permeabilization (LMP) [51] in control treated cells (green arrows in **Fig. 5J**), alongside numerous deacidified lysosomes (black arrows). These morphological changes are characteristic of cells undergoing apoptosis, suggesting that high cytosolic viral loads trigger LMP-mediated cell death pathways to facilitate late-stage viral release [51, 52]. In contrast, SB431542-treated cells maintained acidic lysosomal compartments, correlating with reduced viral titers **(Fig. 5J, right panel and zoomed view)**. To determine whether this inhibitory effect is mediated through canonical TGF-β signalling inhibition in addition to its direct action on ORF3a (off-target effect), we analyzed TGF-β1 gene expression kinetics during infection. As shown in **Fig. 5K**, TGF-β1 expression was significantly upregulated following SARS-CoV-2 infection, consistent with previous reports linking SARS-CoV-2 to TGF-β/Smad3 signaling activation and apoptosis induction in multiple organs [53, 54]. The TGF-β pathway regulates the Coordinated Lysosomal Expression and Regulation (CLEAR) network through transcription factor EB (TFEB). Analysis of CLEAR network genes from our transcriptome data revealed that SB431542 suppressed this gene network at 20 hpi, with a less pronounced effect at 40 hpi **(Fig. 5L).** TFEB transcript levels were significantly reduced at 12 hpi by SB431542 treatment **(Fig. 5M)**, suggesting early inhibition of lysosomal biogenesis. The differential temporal effect of SB431542 on the CLEAR network likely reflects the ability of the compound to counteract ORF3a-mediated dysfunction during early infection, while high viral loads at later timepoints may partially overcome this inhibitory effect.

Since several SARS-CoV-2 proteins including membrane protein, nucleocapsid protein, and ORF3a, are known to induce TGF-β-mediated apoptosis [55, 56], we sought to determine whether apoptosis induction facilitates viral replication. For this, we first analyzed GADD45b expression kinetics, a key positive mediator of TGF-β-induced apoptosis [13]. GADD45b expression was concomitantly upregulated throughout infection but was markedly suppressed by SB431542 treatment **Fig. 5N)**, suggesting that SB431542 inhibits apoptosis via canonical suppression of Smad2/3 signalling and GADD45b expression.

To further characterize the temporal dynamics of apoptosis during infection, we examined expression of BAX, a pro-apoptotic factor regulated by GADD45b [57]. Interestingly, while SB431542 suppressed BAX expression until ∼24 hpi, high viral loads observed at ∼36 hpi appeared to override this inhibitory effect **(Fig. 5O)**. Given that BAX-mediated apoptosis typically requires an induction window of ∼18-24 hours [58], its upregulation during mid-infection likely facilitates virus release through apoptosis during late stages. Transcriptome analysis of genes involved in apoptosis pathway also revealed similar pattern demonstrating that apoptosis signalling was comparatively more active during middle stage of infection and that SB431542 treatment suppressed it **(Fig. 6A)**.

**Figure 6.**
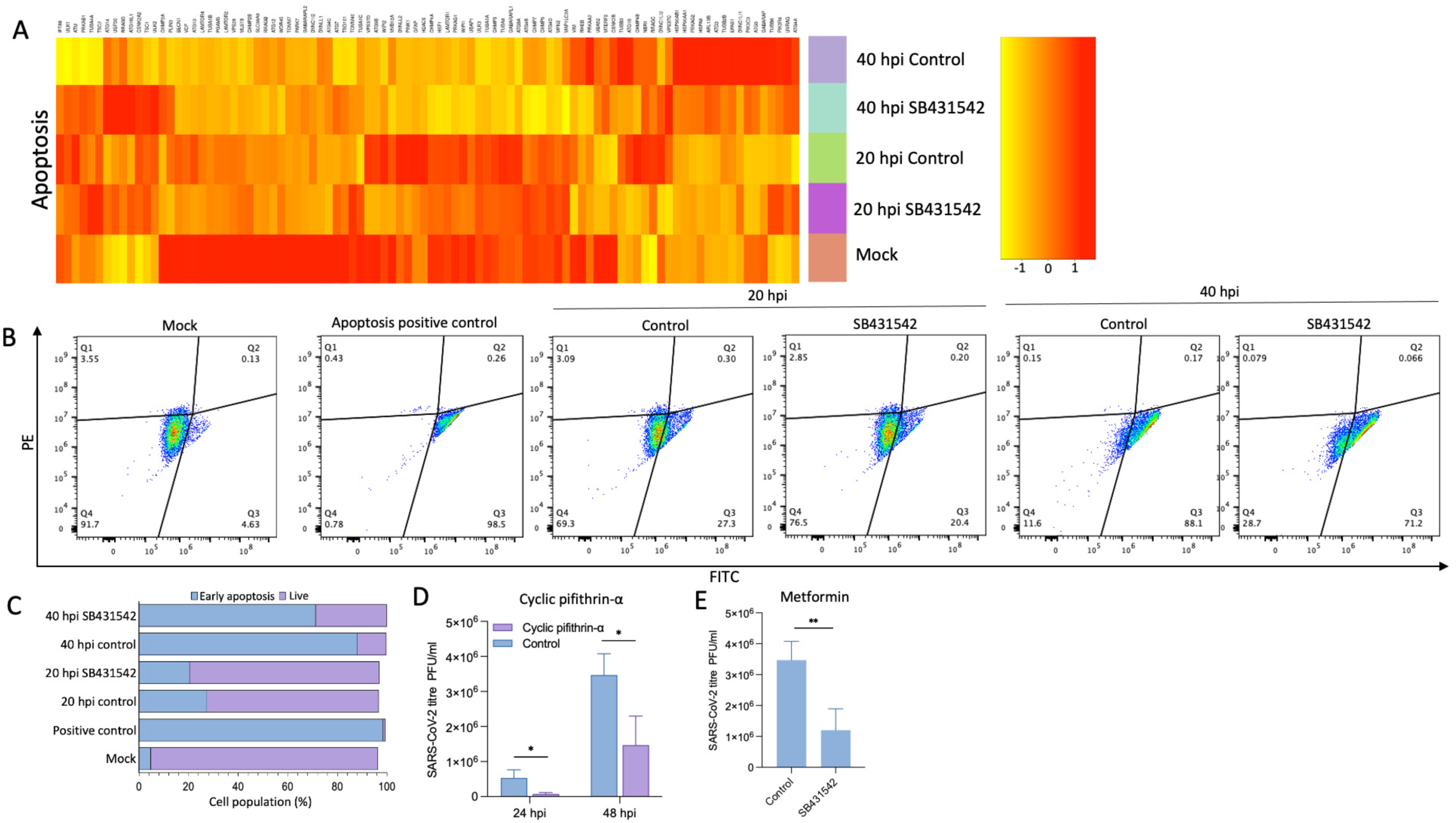
SB431542 blocks SARS-CoV-2 induced apoptosis. **A. RNAseq analysis:** Vero cells were infected with SARS-CoV-2 at 5 MOI for 1 hour followed by treatment with DMSO or SB431542. At 12 hpi, cells were scraped and total RNA was isolated followed by RNAseq analysis as mentioned in methods section. For analysis of apoptosis pathway genes from samples, KEGG pathway filtering was utilized. **B. FACS analysis of early apoptosis:** Vero cells either mock infected or infected with SARS-CoV-2 at 5 MOI, were treated with SB431542 or DMSO. At 20 and 40 hpi cells were trypsinized and treated with JC-1 dye, followed by FACS analysis. Strausoporine treatment group was taken as positive control. **C.** Total cell count as percentage is shown is shown in bar plot **D, E. Antiviral assay:** Vero cells were infected with SARS-CoV-2 at 0.1 MOI for 1 hour followed by treatment with cyclic pifithrin-α **(D),** metformin **(E)** or vehicle controls. For cyclic pifithrin-α, supernatants were harvested at 24 and 48 hpi, and for metformin supernatants were harvested at 48 hpi, followed by virus quantification using plaque assay. Values represent means ± SD from at least three independent experiments. Statistical significance was determined using Student’s t-test (ns = non-significant, *P < 0.05; **P < 0.01).

Next, we quantified apoptotic signatures using JC-1 dye, which measures mitochondrial membrane potential, a key indicator of early apoptosis. Flow cytometric analysis revealed that at 20 hpi, approximately 27.3% of infected cells exhibited early apoptotic signatures, which increased dramatically to 88.1% by 40 hpi **(Fig. 6B)**. SB431542 treatment significantly reduced these apoptotic populations to 20.4% at mid-infection and 71.2% at late stages **(Fig. 6C)**, confirming its time-dependent suppression of virus-induced apoptosis.

To establish a causal relationship between apoptosis and viral egress, we treated infected cells with cyclic pifithrin-α, a selective apoptosis inhibitor [59, 60]. This treatment significantly reduced viral titers at both early (∼24 hpi) and late (∼48 hpi) stages of infection **(Fig. 6D)**, phenocopying the effect of SB431542. Similarly, metformin, which induces autophagy and has been shown to normalize mitochondrial function, also suppressed viral titers at 48 hpi **(Fig. 6E)**.

Collectively, these data demonstrate that SARS-CoV-2 exploits TGF-β-mediated apoptosis as an additional mechanism for efficient virion release during late-stage infection. SB431542 attenuates this process through canonical inhibition of TGF-β/Smad signalling, thereby suppressing the downstream apoptotic cascade. This finding bridges a critical gap between clinical observations of elevated apoptotic cell counts in lung specimens from fatal COVID-19 cases and the molecular basis of late-stage SARS-CoV-2 egress [18–20].

### SB431542 demonstrates *in vivo* efficacy against lethal IBV challenge

To assess the translational potential of SB431542 from *in vitro* to *in vivo* models, we utilized infectious bronchitis virus (IBV), a member of the *Coronaviridae* family that causes lethal pathology in embryonated chicken eggs. Following determination of non-cytotoxic concentrations **(Fig. 7A)**, SB431542 was administered to 10-day-old specific pathogen-free (SPF) embryonated chicken eggs via the allantoic route, concomitant with IBV challenge at 100 EID_50_ (egg infecting dose). SB431542 conferred robust, dose-dependent protection against lethal IBV infection, as evidenced by significantly improved survival rates compared to vehicle-treated controls **(Fig. 7B)**.

**Figure 7.**
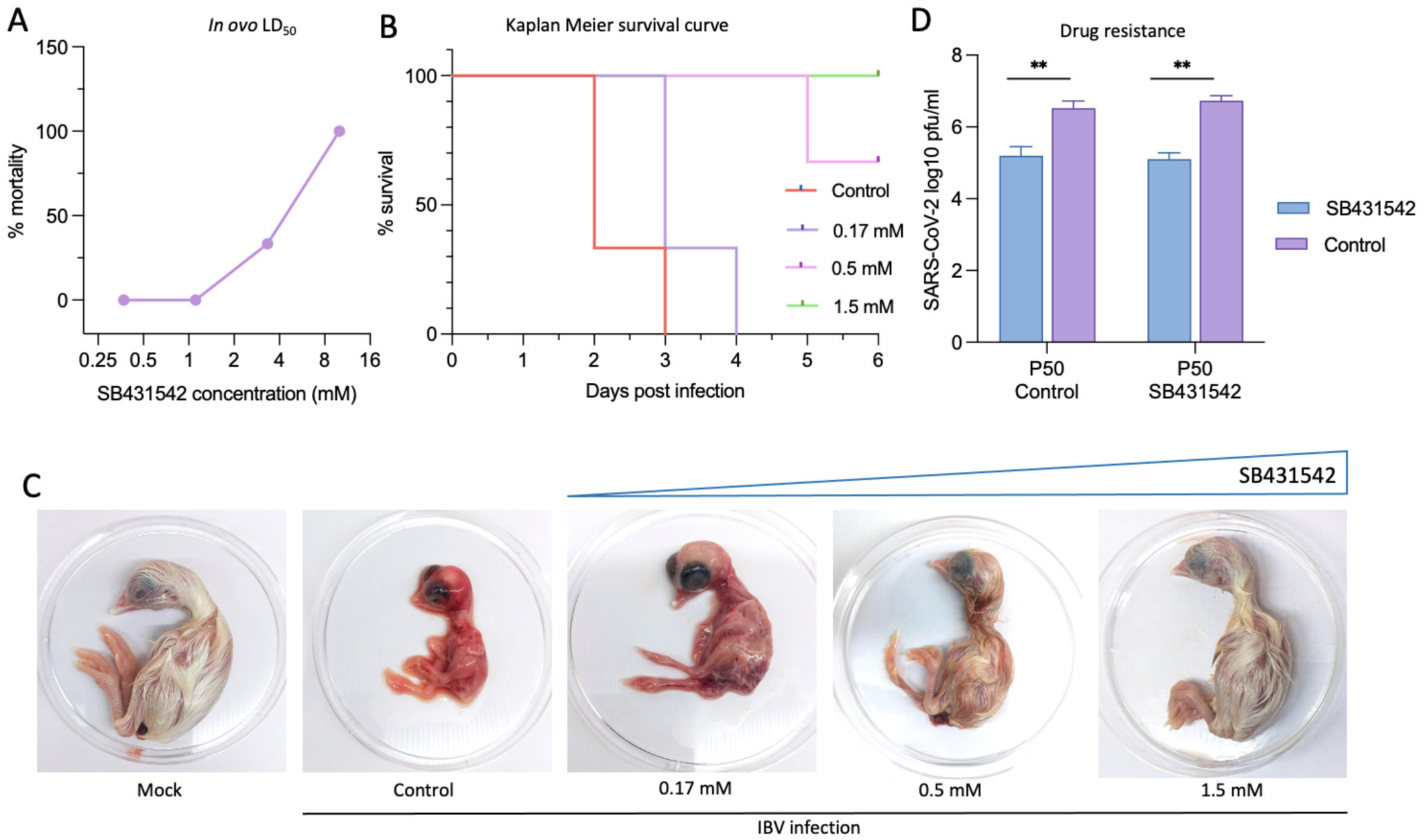
*In ovo* antiviral efficacy and drug resistance study. **A. Determination of LD_50_.** Indicated concentrations of SB431542, in triplicates, were inoculated in 10 day old embryonated SPF chicken eggs. The viability of the embryos was examined for up to 5 days post inoculation and the LD_50_ was determined to be 7.60 mM (Reed-Muench method). . **B. Survival curve.** Three-fold serial dilutions of SB4531542 or DMSO were inoculated, in triplicates, in 10 days old SPF embryonated eggs via the allantoic route, followed by infection with IBV at 100 EID_50_. The viability of the eggs was examined for six days post-infection. The EC_50_ was calculated to be 0.22 mM. **C. Morphological changes:** Morphological changes in the chicken embryos at different drug regimens following IBV challenge is shown. **D. Evaluation of SARS-CoV-2 resistance against SB431542:** ARS-CoV-2 was sequentially passaged 50 times in Vero cells in the presence of SB431542 or DMSO. For each passage, confluent monolayers of Vero cells were infected with SARS-CoV-2 at an MOI of 0.1, followed by washing with PBS and the addition of fresh DMEM supplemented with SB431542 (0.25 uM) or 0.05% DMSO. At the end of P50, the fitness of SB431542- or DMSO-passaged viruses was evaluated again against SB431542. Values represent means ± SD from at least three independent experiments. Statistical significance was determined using Student’s t-test (**P < 0.01).

Morphological assessment further corroborated these findings, revealing that SB431542 treatment dose-dependently prevented the characteristic stunted embryonic development associated with lethal IBV infection **(Fig. 7C)**. Pharmacological evaluation demonstrated that SB431542 exhibited a lethal dose 50 (LD_50_) of 7.60 mM and an effective concentration (EC_50_) of 0.22 mM, yielding a therapeutic index (TI = LD_50_/EC_50_) of 34.54, indicating a substantial safety margin for potential therapeutic applications. These results suggest that SB431542 effectively mitigates IBV-induced lethality in embryonated chicken eggs and underscores its potential as a broad-spectrum antiviral agent.

### Long-term selection pressure of SB431542 does not produce viral escape mutants

To address the critical concern of resistance emergence that plagues many direct-acting antivirals, we conducted prolonged selection pressure experiments by sequentially passaging SARS-CoV-2 in Vero cells for 50 generations in the presence of either SB431542 or vehicle control (DMSO). Remarkably, following this extensive passaging regimen, both control-passaged (P50 control) and drug-passaged (P50 SB431542) viral variants remained equally susceptible to SB431542 treatment **(Fig. 7D).** This exceptional genetic barrier to resistance likely stems from SB431542’s multifaceted mechanism of action, simultaneously targeting both host factors (TGF-β signalling, apoptosis) and viral component (ORF3a), thereby imposing multiple selective constraints that collectively impede the emergence of viable escape mutants. These findings starkly contrast with the rapid resistance development observed with direct-acting antivirals such as remdesivir or monoclonal antibodies, which frequently drive the evolution of escape mutants under selection pressure [61–63].

Collectively, these results highlight the potential utility of SB431542 as a robust therapeutic candidate with minimal risk of resistance development, addressing one of the most significant challenges in antiviral drug development[64, 65].

## Discussion

Drug repurposing, particularly of host-directed agents (HDAs), represents a compelling antiviral strategy by establishing high genetic barriers against viral escape mutations while offering broad-spectrum activity [66, 67]. Unlike direct-acting antivirals (DAAs), which often lead to rapid resistance development [68], numerous HDAs have been successfully repurposed as antiviral drugs or are undergoing preclinical and clinical evaluation [66, 67].

Our comprehensive investigation reveals that an established TGF-β inhibitor SB431542 (an HDA) exhibits potent antiviral activity against coronaviruses through an unprecedented multiple action mechanism: inhibition of viral entry via canonical TGF-β/Smad signaling modulation, direct binding to viral ORF3a protein which disrupts lysosomal function, and attenuation of TGF-β-induced apoptosis that facilitates late-stage viral release.

Our time-of-addition and step-specific assay revealed that SB431542 suppressed maximum SARS-CoV-2 replication when administered either before infection or within early post-infection periods. While its inhibitory effect on viral entry through TGF-β/Smad-mediated modulation of furin expression was recently reported previously [8], our study uncovered two additional mechanisms targeting the later stages of the viral life cycle. Notably, despite comparable viral protein synthesis, we observed a significant reduction in both intracellular and extracellular virions following SB431542 treatment. This indicates potential interference with viral assembly and egress pathways rather than replication machinery.

The observed disruption of nucleocapsid-RNA association in RNA-IP assays provided the first mechanistic insight into how SB431542 impairs virion assembly **(Fig. 4C)**. Given the established role of ORF3a in lysosomal dysfunction and autophagy subversion during SARS-CoV-2 infection, we hypothesized that SB431542 might directly target this viral protein. Our *in silico* analyses revealed that SB431542 binds to ORF3a with high affinity (ΔG = -13.32 kcal/mol), outcompeting bacteriochlorin (ΔG = -5.28 kcal/mol) at a site overlapping with the Saposin-A interaction interface [37]. This binding site is critically positioned to disrupt the canonical role of ORF3a in inhibiting autophagosome-lysosome fusion and promoting lysosomal deacidification, the processes essential for preserving virion integrity during egress [37].

Immunofluorescence studies also validated this mechanism, demonstrating that SB431542 treatment normalized LC3-GFP distribution patterns, restored perinuclear LAMP-1 localization, and reacidified lysosomes in infected cells **(Fig. 2)**. These effects collectively reversed the lysosomal dysfunction induced by SARS-CoV-2 infection, consistent with the predicted interaction of the compound with ORF3a. The relevance of this pathway was further confirmed when metformin, a known autophagy inducer, similarly suppressed viral titers **(Fig. 5D)** [45]. These findings complement recent studies identifying lysosomal exocytosis as a distinctive egress mechanism for β-coronaviruses, and suggest that therapeutic modulation of this pathway represents a viable antiviral strategy[49].

A particularly novel aspect of our study is the characterization of temporal dynamics in SARS-CoV-2 egress mechanisms. While previous investigations have primarily focused on early infection stages (≤24 hpi) [2, 49, 50], our one-step growth curve analysis revealed a substantial 340% increase in viral titers between 36-48 hpi. Coupled with the emergence of cytopathic effects only after ∼ 42 hpi, this temporal progression suggested potentially additional viral egress strategies during prolonged infection. Indeed, our JC-1 staining experiments demonstrated progressive induction of apoptosis, with early apoptotic signatures increasing from 27.3% at 20 hpi to 88.1% by 40 hpi. Mechanistically, we observed time-dependent upregulation of TGF-β1 and its downstream effector GADD45b during infection, with subsequent induction of the pro-apoptotic factor BAX **(Fig. 5I, 5J and 6A)** [57].

The relationship between apoptosis and late-stage viral release was confirmed when cyclic pifithrin-α, a selective apoptosis inhibitor [60], significantly reduced viral titers **(Fig. 6D)**. SB431542 treatment similarly suppressed apoptotic signatures and reduced viral titers, indicating that inhibition of TGF-β-mediated apoptosis constitutes a third mechanism through which this compound exerts antiviral effects. This finding bridges an important gap between the clinical observation of elevated apoptotic cell counts in lung specimens from fatal COVID-19 cases [16, 18–20], and the molecular basis of late-stage SARS-CoV-2 egress, suggesting that increasing viral load-induced apoptosis serves as a release mechanism complementary to lysosomal exocytosis during sustained infection.

Furthermore, the translational potential of SB431542 was validated in an IBV infection model using embryonated chicken eggs, where dose-dependent protection against lethal viral challenge was observed **(Fig. 7C)**. The calculated therapeutic index of 34.54 highlights a favourable safety margin, although further studies in mammalian models will be necessary to fully evaluate its pharmacological profile. Perhaps most significantly, sequential passaging of SARS-CoV-2 for 50 generations under SB431542 selection pressure failed to generate resistant variants **(Fig. 7D)**. This high genetic barrier to resistance likely stems from SB431542’s multifaceted mechanism of action targeting both host factors (TGF-β signaling, apoptosis) and viral component (ORF3a). This contrasts with direct-acting antivirals such as remdesivir or monoclonal antibodies, which frequently drive the evolution of escape mutants under selection pressure [61–63].

Collectively, our findings establish SB431542 as a promising antiviral candidate with a remarkable EC_50_ of 515 nM against SARS-CoV-2, significantly more potent than the FDA-approved remdesivir (EC_50_: 1.65 μM) in comparable Vero cell models [24]. The compound’s unique triple-mechanism approach – inhibiting viral entry via TGF-β/Smad signaling modulation, disrupting ORF3a-mediated lysosomal dysfunction affecting assembly and egress, and attenuating TGF-β-induced apoptosis during late-stage infection provides multiple barriers against viral replication while minimizing resistance development.

However, certain limitations warrant further investigation. While our study demonstrates efficacy *in vitro* and *in vivo* using embryonated chicken eggs as a model system, additional preclinical studies using mammalian models are necessary to evaluate pharmacokinetics, toxicity profiles, and efficacy against emerging SARS-CoV-2 variants. Additionally, comprehensive toxicological studies will be needed to assess potential side effects associated with systemic TGF-β inhibition, particularly in the context of prolonged treatment.

## Author contributions

Conceptualization, N.Ku., A.V. Methodology and investigation, A.V., G.K., N.Kh. Cloning and Biochemical assays, A.V., G.K., N.Kh., Y.C., A.K., *in silico* work, A.V., B.E.M., *in ovo* work, A.V., J.R., H.K., S.D., Data analysis, R.K., R.T., S.S., N.Ku. Supervision, N.Ku. Writing-original draft, A.V., Writing-review and editing, A.V., N.Ku.

## Declaration of Competing Interest

The authors declare that they have no known competing financial interests or personal relationships that could have appeared to influence the work reported in this paper.

## Disclosure statement

The authors declare that the work was conducted in the absence of any commercial or financial relationships that could be construed as a potential conflict of interest.

## Acknowledgements

This work was supported by Science and Engineering Research Board (SERB), Department of Science and Technology, Government of India (Grant Number CVD/2020/000103 to N. Ku.). The funders had no role in study design, data collection and analysis, decision to publish, or preparation of the manuscript.

## Materials and methods

### Cells and Viruses

Vero cells (African green monkey kidney cells) were cultured in Dulbecco’s Modified Eagle Medium (DMEM) supplemented with 10% fetal bovine serum (FBS) and antibiotics. SARS-CoV-2 (SARS-CoV-2/Human-tc/India/2020/Hisar-4907) was propagated in Vero cells under Biosafety Level 3 (BSL-3) conditions at the National Centre for Veterinary Type Cultures (NCVTC), ICAR-National Research Centre on Equines, Hisar, India. Viral titers were quantified as plaque-forming units per ml (PFU/mL) by plaque assay.

For *in vivo* studies, infectious bronchitis virus (IBV), a member of the *Coronaviridae* family, was used to infect specific pathogen-free (SPF) embryonated chicken eggs. IBV was propagated and quantified as egg infectious dose 50 (EID_50_) per ml.

### Inhibitors

SB431542 was purchased from Sigma-Aldrich and dissolved in dimethyl sulfoxide (DMSO). Bafflomycin, cyclic pifithrin-α, metformin and strausoporine were procured from MedChemExpress.

### Cytotoxicity Assay

The cytotoxicity of drugs in Vero cells was determined using the MTT assay. Briefly, Vero cells were seeded into 96-well plates and treated with serial dilutions of inhibitors for 96 hours. Cell viability was measured by adding MTT reagent (5 mg/mL) to each well, followed by incubation at 37°C for 4 hours. The formazan crystals formed were dissolved in DMSO, and absorbance was measured at 570 nm using a microplate reader. The half-maximal cytotoxic concentration (CC_50_) was calculated using Reed-Muench method.

### Plaque Assay

Plaque assays were performed to quantify SARS-CoV-2 titers. Confluent monolayers of Vero cells in 12-well plates were infected with serial dilutions of virus-containing supernatants for 1 hour at 37°C. After infection, supernatants containing viruses were removed and cells were overlaid with L-15 media containing 2% agarose and incubated for 96 hours at 37°C. Plaques were visualized by staining with crystal violet solution and counted to determine viral titers.

### Time-of-Addition Assay

Confluent monolayers of vero cells were infected with SARS-CoV-2 at an MOI of 5, followed by washing with PBS for 5 times, supplemented with fresh DMEM and treated with SB431542 or vehicle control (DMSO) at various time points post-infection (-0.5 h to 36 hpi). Supernatants were collected at 48 hpi, and viral titers were quantified by plaque assay.

### Virus step-specific assays

For attachment assay, Vero cell monolayers were pre-treated with SB431542 or vehicle control for 1 hour, followed by SARS-CoV-2 infection (MOI 5) at 4°C for 1 hour. After five PBS washes to remove unbound virus, cell lysates were prepared by freeze-thaw cycling and viral titers were quantified by plaque assay.

For virus entry studies, prechilled Vero cell monolayers were infected with SARS-CoV-2 (MOI 5) at 4°C for 1 hour to permit attachment. Following five PBS washes, cells were incubated with SB431542 or vehicle control at 37°C for 1 hour to allow virus entry. After removing extracellular virus by PBS washing, cells were maintained in inhibitor-free DMEM. Infectious virus particles in supernatants were quantified by plaque assay at 12 hpi. To assess virus release, SARS-CoV-2-infected Vero cells (MOI 5) were incubated until 8 hpi. Cells were then washed and treated with SB431542 or vehicle control. Supernatants were collected at 4 hours post-treatment for plaque assay quantification.

For viral protein synthesis analysis, SARS-CoV-2-infected (MOI 5) or mock-infected Vero cells were treated with SB431542 or vehicle control at 3 hpi. Cell lysates were prepared at 12 hours post-infection for western blot analysis of viral and cellular proteins.

To quantify viral RNA/DNA, SARS-CoV-2-infected Vero cells (MOI 5) were treated with SB431542 or vehicle control at 2 hours post-infection. At 12 hours post-infection, total RNA/DNA was extracted. cDNA was synthesized using oligo dT primers (Fermentas, Hanover, USA) for mRNA quantification. SARS-CoV-2 *N* gene and β-actin expression were quantified by qRT-PCR as described previously[69].

### Lysosomal Function Assays

To assess lysosomal function, Lysotracker Red DND-99 dye (Thermo Fisher Scientific) and Hoechst dye were used to stain acidic lysosomes and nucleus respectively in infected Vero cells treated with SB431542 or vehicle control. Cells were incubated with the dye for 30 minutes at 37°C according to manufacturer’s protocol, washed with PBS, and visualized under a fluorescence microscope.

For transcriptomic analysis of autophagy-related genes, RNA was extracted from infected cells using TRIzol reagent (Thermo Fisher Scientific). cDNA synthesis was performed using oligo-dT primers, followed by quantitative real-time PCR (qRT-PCR) using SYBR Green Master Mix (Bio-Rad). Gene expression levels were normalized to β-actin as a housekeeping gene.

### Apoptosis Assays

Mitochondrial membrane potential was assessed using JC-1 dye (Thermo Fisher Scientific). Infected Vero cells treated with SB431542 or vehicle control were harvested at mid-infection (20 hpi) and late stages (40 hpi). Cells were stained with JC-1 dye according to the manufacturer’s protocol and analyzed using a BD FACSARIA flow cytometer. The percentage of early apoptotic cells was determined based on red-to-green fluorescence ratios. Strausoporine treated cells were taken as positive control.

### RNA Immunoprecipitation Assay

RNA immunoprecipitation (RNA-IP) assays were performed to evaluate the interaction between viral RNA and nucleocapsid protein. Infected Vero cells treated with SB431542 or vehicle control were crosslinked with formaldehyde to stabilize RNA-protein interactions at 12 hpi by incubating with 1% formaldehyde for 10 minutes and quenched with 125 mM glycine. Cells were lysed in immunoprecipitation/lysis buffer (150 mM NaCl, 50 mM Tris-HCl [pH 7.5], 5 mM EDTA, 0.5% NP-40, 1% Triton X-100, and protease/phosphatase inhibitor cocktail) by incubating for 30 mins followed by sonication (Qsonica Q500) using six 15-second pulses at 40% amplitude, clarified by centrifugation (12,000 g, 10 minutes), and supplemented with 10 units of RiboLock RNase Inhibitor (Invitrogen, Carlsbad, USA). Cell lysates were immunoprecipitated using SARS-CoV-2 anti-nucleocapsid antibody conjugated to Protein A Sepharose beads (Sigma-Aldrich). α-MNK1 antibody and IP buffer served as nonreactive and beads controls, respectively. Following five washes with IP buffer, RNA-protein complexes were reverse crosslinked, and RNA was extracted using TRIzol reagent for downstream qRT-PCR analysis. Values were normalized to input RNA. All experiments were performed in triplicate.

### Cloning and overexpression of ORF3a

SARS-CoV-2 ORF3a was cloned into pDONR221 vector and subsequently transferred to pDEST40 vector using GATEWAY cloning technology (Invitrogen, Carlsbad, CA, USA) according to manufacturer’s instructions. The ORF3a was amplified with attB-flanked primers (forward primer 5’-GGGGACAAGTTTGTACAAAAAAGCAGGCTTCACCATGGATTTG TTTATGAGAATCTTCACAATTGG-3’ and reverse primer 5’-GGGGACCACTTTGTACAAGAAAGCTGGGTGCAAAGGCACGCTAGTAGTCGTC-3’). Cloned constructs were verified by sangar sequencing.

### Immunofluorescence studies

HeLa cells grown in 4-chambered slides at 5% confluency in DMEM supplemented with FBS and were transfected with 1 μg of LC3-GFP (addgene #11546) and pDEST40:ORF3a or empty plasmid using Lipofectamine 3000. At 24 hours post-transfection (hpt), cells were treated with SB431542 or vehicle control and at 48 hpt cells were subjected to immunofluorescence studies for visualization of LC3-GFP puncta. In case of transfection in Vero cell with LC3-GFP, at 48 hours post-transfection SARS-CoV-2 infection was given at 5 MOI in the presence of SB431542 or vehicle control and at 12 hpi cells were subjected to immunofluorescence studies as described previously [70].

### *In Vivo* Antiviral Efficacy in Embryonated Chicken Eggs

SPF embryonated chicken eggs (10 days old) were procured from Indovax Pvt Ltd., Hisar, India. To determine the lethal dose 50 (LD_50_) of SB431542, eggs were inoculated via the allantoic route with serial dilutions of SB431542 or DMSO as a vehicle control. Egg viability was monitored daily for up to six days post-inoculation.

For antiviral efficacy studies, eggs were infected with IBV at an EID of 100 via the allantoic route and treated with SB431542 or DMSO immediately post-infection. Eggs were monitored for survival and foetal growth for up to six days post-infection.

### RNA extraction, library preparation, and Illumina sequencing

Vero cells (80-85% confluent) were infected with SARS-CoV-2 (MOI 5), treated with SB431542 at 1 hour post-infection, and harvested at 20 and 40 hpi for total RNA extraction using TRIzol reagent (Takara, China).

Poly(A) mRNA was enriched from total RNA (500 ng) using the NEBNext® Poly(A) mRNA Magnetic Isolation Module (New England Biolabs, MA, USA) according to the manufacturer’s protocol. Libraries were prepared using the NEBNext® Ultra™ II RNA Library Prep Kit. Enriched mRNA was fragmented in magnesium-based buffer at 94°C for 10 minutes using NEBNext® Random Primers to generate ∼300 nucleotide inserts. The fragmented RNA was reverse transcribed into first-strand cDNA and was followed by double-stranded DNA conversion and purification using 1.8X AMPure XP beads (Beckman Coulter, CA, USA). The purified dsDNA was subjected to end repair, 3’ adenylation, and loop adapter ligation.

Following USER enzyme treatment, adapter-ligated products were size-selected using AMPure XP beads to obtain 400-600 bp libraries. Libraries were amplified through 12 PCR cycles using NEBNext® Ultra II Q5 Master Mix and Multiplex Oligos, then purified with 0.9X AMPure XP beads and eluted in 15 µl of 0.1X TE buffer.

Adapter ligated products were treated with USER enzyme and size-selected using AMPure XP beads (Beckman Coulter, CA, USA) to target a library size of 400–600 bp. The cDNA libraries were amplified through 12 PCR cycles using NEBNext® Ultra II Q5 Master Mix and NEBNext® Multiplex Oligos for Illumina. The amplified libraries were purified with 0.9X AMPure XP beads and eluted in 15 µl of 0.1X TE buffer.

Library quality was assessed using a Qubit Fluorometer (Invitrogen, Life Technologies, USA) for quantification and Tapestation system (Agilent Technologies, USA) with HSDNA kit for size distribution analysis. Pooled libraries underwent cluster generation on the c-Bot system followed by paired-end sequencing (2×150 bp) on an Illumina HiSeq X10. Demultiplexing and adapter trimming were performed using CASAVA v1.8.2 (Illumina Inc.).

### RNA-seq data analysis

Raw read quality was assessed using FastQC (v0.11.8) to examine base quality score distribution, sequence quality score distribution, average base content per read, and GC content distribution. Adapter sequences (AGATCGGAAGAGC) were removed using Trim Galore (v0.6.2), which also performed automated quality trimming. Reads shorter than 20 bp or with low-quality ends (Phred score <20) were discarded. The *Chlorocebus sabaeus* reference genome (Ensembl release 110) and SARS-CoV-2 genome were indexed using BWA (v0.7.17), and pre-processed reads were aligned using BWA-MEM with default parameters. Mapped reads were quantified at the gene level using Samtools (v0.1.19).

Differential expression analysis was performed using DESeq (v1) to identify genes with significant expression changes. Differentially expressed genes (DEGs) were defined using threshold criteria of fold change ≥2 and p-value <0.05. DEGs were functionally annotated using BLASTx (v2.2.29+) against the NR database, with annotations retrieved from UniProt and KEGG databases. Gene Ontology (GO) terms were visualized using WEGO with a log10-scaled y-axis. Expression patterns were visualized through volcano plots and MA plots generated using custom R scripts, while heatmaps were created using MeV software (v4.8.1).

### Generation of protein-ligand complexes

To investigate the binding interactions of SB431542 with SARS-CoV-2 ORF3a, protein-ligand complexes were generated using the diffusion-based structure prediction tool Boltz [32]. This approach leverages machine learning models trained on datasets of bound complexes by simulating "diffusion" from a disordered state to a bound state. The protein template was based on the experimentally determined cryo-EM structure of ORF3a dimer (PDB ID: 6XDC, resolution 2.9 Å)[71]. To maintain the physiologically relevant dimeric state of ORF3a, structure prediction was performed with two copies of both the protein sequence and ligand SMILES string. For analysis of single ligand binding (SB431542), the second ligand was subsequently removed from the complex. A total of 25 structural models were generated and as a reference compound, we modelled bacteriochlorin binding to ORF3a using identical protocols, as bacteriochlorin is structurally similar to chlorin compounds previously reported to interact with the chloride binding site of ORF3a[31, 72].

The Boltz-generated complexes were further refined to ensure compatibility with downstream molecular dynamics simulations. This approach was chosen over traditional docking methods (e.g., GLIDE, AutoDock, and RosettaDock) due to their limitations in accurately predicting binding affinities when experimental data is unavailable. Diffusion-based modeling provides an alternative capable of generating high-confidence protein-ligand complexes with performance comparable to AlphaFold3[73].

### Molecular dynamics simulation protocol

Molecular dynamics (MD) simulations were performed using GROMACS with the CHARMM36 force field for proteins[74]. For small molecule parameterization, we employed espaloma, a machine learning-based approach that generates classical force field parameters[75]. The all-atom simulations followed an adjusted protocol based on established methodologies[76, 77].

The prameters can be found in the following GitHub repository (<u>link</u>). The protocol began with a series of minimization steps in vacuum, consisting of steepest descent minimization, followed by conjugate gradient minimization (repeated twice), and a single minimization using the LBFGS method, all implemented within GROMACS. After these initial minimization steps, the protein-ligand complex was solvated and neutralized with ions, and the same series of minimization procedures was repeated for the solvated system.

System equilibration was conducted in two phases: an initial 2 ns simulation followed by a 0.5 ns relaxation simulation, both performed in the NPT ensemble using scripts 3-pr-npt.mdp and 4-npt-relax.mdp, respectively. Production simulations were then carried out for 500 ns in the NPT ensemble. All simulations were performed at a temperature of 300K and pressure of 1 bar.

For trajectory analysis, we calculated several standard metrics including root mean square deviation (RMSD) to assess overall structural stability, radius of gyration (Rg) to detect potential protein unfolding, protein-ligand contacts (using a 4 Å distance cut-off), and root mean square fluctuation (RMSF) to evaluate residue mobility. All analyses were performed using custom Python scripts with the biotite library. Simulation protocols and scripts are available at the GitHub repository (<u>link</u>).

### Binding free energy calculations

Multiple computational approaches were employed to estimate binding affinities. First, MM-GBSA calculations were performed on the last 1000 frames of each simulation using gmx_MMPBSA [34, 35]. The implicit solvent model was employed with a salt concentration of 0.15 M. Next, non-equilibrium pulling simulations were conducted by applying a harmonic restraint between the centre of mass of the ligand and a reference point outside the binding pocket[78]. The restraint was moved at a constant velocity of 10 nm/ns over 2 ns with a force constant of 1000kJ/mol/nm². Only a single pulling simulation was performed as preparation step for the subsequent umbrella run. And finally more accurate binding free energy estimation, we used umbrella sampling on frames selected from the pulling trajectory [36]. Initially a total of 20 windows with equidistance spacing over the generated pulling trajectory where chosen. After evaluation of the resulting histograms windows were added manually at frames best representing the center of gaps inbetween histograms. A total of 45 windows for SB431542 and 42 windows for Bacteriochlroin were simulated using this approach. For an exact documentation as well as result of the performed umbrella sampling runs see the GitHub repository (<u>link</u>). Each window was sampled for 10 ns with a restraint force constant of 5 kcal/mol/Å². The weighted histogram analysis method (WHAM) was used to reconstruct the potential of mean force (PMF) and calculate binding free energies (ΔG). The binding free energy was estimated by calculating the difference between PMF values of bound and unbound states, providing a more thermodynamically rigorous assessment of ligand affinity.

All trajectory analyses including root mean square deviation (RMSD), root mean square fluctuation (RMSF), radius of gyration (Rg), and protein-ligand contacts were performed using a customized Python scripts utilizing the Biotite library.

### Statistical analysis

All experiments were conducted in triplicates unless otherwise specified. Data are presented as mean ± standard deviation (SD). Statistical significance between groups was determined using Student’s *t*-test (*p* < 0.05 considered significant).

### SB431542 suppresses SARS-CoV-2 release from Vero cells during late hours of infection

